# HLA-G engineering reprograms CAR-T cells with an immune privilege

**DOI:** 10.64898/2026.05.10.723228

**Authors:** Yuanyuan Xie, Shangqing Hong, Chenling Xu

## Abstract

Personalized T cell therapy empowered by chimeric antigen receptor (CAR) that recognizes specific tumor antigen has cured numerous blood cancer patients since its initial approval in 2017. However, its access to a broader population has been limited by the unavailability of an off-the-shelf product derived from an allogeneic donor that can evade immune rejection, which is mediated by polymorphic class I and class II human leukocyte antigens (HLAs). Since class II HLAs are only expressed in specialized antigen-presenting cells but not T cells, it might suffice to evade T cells by deleting the common class I HLA light chain Beta-2 Microglobulin (B2M) (*1*). However, *B2M*-deficient cells can trigger a “missing-self” response to activate natural killer (NK) cells (*2*), a second function that was evolved to compensate loss of T cell response. Inserting a less polymorphic class I HLA gene encoding a known NK inhibitory ligand, namely HLA-E or HLA-G (*3*), into the *B2M* locus so that the endogenous *B2M* expression is disrupted could theoretically allow evasion of both T and NK cells. Despite being a seemingly better candidate in that HLA-G is uniquely expressed in immune-privileged sites such as the placenta with a believed function in protecting the fetus from immune rejection by the pregnant mother, whereas ubiquitously-expressing HLA-E is known to bind both inhibitory and activating NK receptors (*4*, *5*), only HLA-E engineering has been attempted yet without convincing success *in vivo* (*6*, *7*). Here, we generate an off-the-shelf CAR-T product with *B2M* replaced by a gene fusion encoding an HLA-G single-chain trimer under minimally impacted *B2M* epigenetic landscape, and observe its immune evasion property and a tumor-inhibitory function that is equivalent to its autologous control using a humanized mouse model for the first time with T and NK cells reconstituted from a donor with a distant HLA haplotype. HLA-G engineering may thus reprogram T cells into an immune-privileged state that can be utilized for all cell-based therapies.

## Results

A landmark paper was published in 2017 by the Sadelain group demonstrating that inserting a CAR into the T cell receptor α constant (*TRAC*) locus to drive CAR expression under the endogenous epigenetic landscape of *TRAC* gives rise to more persistent CAR-T cells compared to the traditional retroviral transduction method (*8*). Notably, this phenotype was only revealed in *in vivo* rather than *in vitro* studies (*8*), consistent with unique upstream epigenetic modulators contributed from an *in vivo* microenvironment, which may apply to our study as well. Using the same guide RNA (gRNA) from this paper (*8*), we firstly set to knock in a clinically validated CD19-CAR (*9*) by CRISPR/Cas9 into the *TRAC* locus while knocking out *TRAC* expression to generate *TRAC^CD19-CAR/CD19-CAR^* (YP001) cells (Fig 1a, S1a and Table S1). These cells should not cause graft-vs-host disease (GvHD) in the recipient due to the *TRAC* knockout, a key safety feature for an off-the-shelf CAR-T product. We then set to generate *TRAC^CD19-CAR/CD19-CAR^;B2M^B2M-HLA-G/B2M-HLA-G^*(YP103) using a double knockin/knockout strategy where a single-chain trimer of HLA-G fused to B2M and an HLA-G signal peptide (*10*) was also inserted into the *B2M* locus (Fig 1a, S1a and Table S1). Since similar HLA-E-engineered products were previously reported by other groups (*6*, *11*), we also planned to make the HLA-E version (YP102) for comparison (Fig S1a and Table S1). We set to perform a tumor-bearing xenograft experiment in which allogeneic CAR-T cells can kill tumor cells in a humanized mouse model at a rate impacted by host T and NK cells derived from a distant peripheral blood mononuclear cell (PBMC) donor. Using this model, we tested for similar tumor-inhibitory effects with our allogeneic product and autologous control cells, and for rejection of allogeneic control cells *TRAC^CD19-CAR/CD19-CAR^* (YP001) and *TRAC^CD19-CAR/CD19-CAR^;B2M^-/-^*(YP100, see Fig S1a). Since editing efficiency might differ between single and double knockin experiments, we decided in the beginning to sort cells using a CD19-biotin antibody and use a targeting construct lacking the B2M signal peptide (YP102ΔS, see Fig S1a) as controls for surface expression.

**Fig 1.**
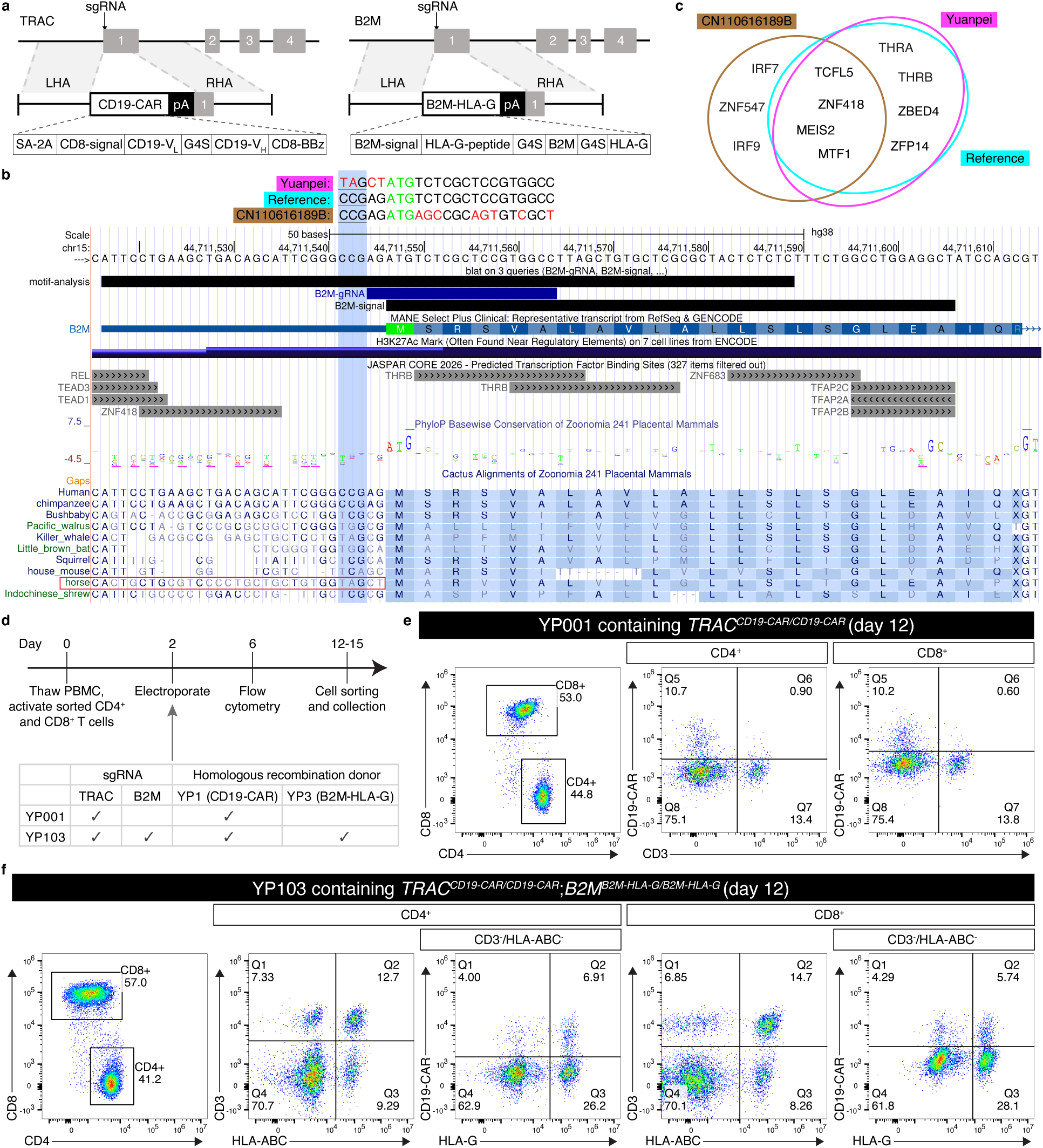
Generation of HLA-G-engineered chimeric antigen receptor T (CAR-T) cells. **a**, Schematic of the genome editing strategy. The CD19-CAR and B2M-HLA-G fusion genes followed by a polyA (pA) sequence are flanked by homology arms (left: LHA; right: RHA) for homologous recombination (HR) mediated integration into the *TRAC* and *B2M* loci, respectively, using CRISPR/Cas9 with single guide RNA (sgRNA) targeting exon 1. Major structural components are illustrated in the fusion gene schematic (also see Table S1). Note that the G4S linker is present in the CD19-CAR fusion gene in between the variable light (CD19-V_L_) and heavy (CD19-V_H_) domains of the single-chain variable fragment (ScFv). **b**, UCSC Genome Browser view underneath three 23 base-pair (bp) sequences (3 bp PAM underlined with blue background plus 20 bp spacer) with red letters indicating mutations compared to the middle reference and green “ATG” letters matching the *B2M* translation initiation site. The horse orthologous non-coding sequence is highlighted with a red square. **c**, Transcription factors that are predicted to bind the three 73 bp sequences as illustrated in the motif-analysis (using FIMO with a default 1×10^-4^ *P* value cutoff) BLAT track (*29*) containing three 23 bp sequences shown in **b**. **d**, Timeline of genome editing (upper) with materials illustrated for generating different CAR-T cells (lower). **e**, **f**, Flow cytometric analysis for YP001 (**e**) and YP103 (**f**) collected on day 12 (see **d**) derived from peripheral blood mononuclear cell (PBMC) donor 416C. See also Fig S1.

Integrated B2M-HLA fusion gene is prone to Cas9 recutting unless enough mismatches are designed in the region of the homologous recombination (HR) donor that is targeted by the B2M gRNA. We used a gRNA described previously by the Sadelain group (*8*) which targets around the B2M translation initiation site and an intuitive way to avoid Cas9 recutting would be to make some synonymous mutations in the gRNA region as done in the patent CN110616189B (Fig 1b). However, we identified two predicted THRB motifs using the UCSC Genome Browser (*12*) JASPAR track (Fig 1b), suggesting that mutations in the corresponding spacer region of the B2M gRNA may disrupt THRB binding and thus the B2M endogenous epigenetic environment. Therefore, we instead utilized alignment tracks to borrow a DNA sequence from a placental mammal ortholog where the NGG PAM region is mutated to block Cas9 binding (Fig 1b). Given that *Streptococcus pyogenes* Cas9 (SpCas9) is known to also bind to non-NGG sequences, such as NGA and NAG (*13*), and DNA sequences in the spacer region that are closer to the PAM are more critical for gRNA interaction (*14*), we selected the sequence from the horse ortholog with four mutations (<u>CCG</u>AG to <u>TAG</u>CT, PAM underlined) (Fig 1b), reasoning that the corresponding epigenetic axis may be evolutionarily conserved. To test this idea, we used the FIMO program (*15*) to predict transcription factor binding sites against the JASPAR motif database (*16*), and found our picked sequence showed the exact same predicted motifs as the reference, whereas the sequence used in CN110616189B added three different motifs at the expense of four existing ones including THRB (Fig 1c). We included these four mutations in the 1 kb homology arms (*17*) flanking the B2M-HLA-E fusion gene with its translated sequence matching 100% to the patent US11813318B2 (Table S1). We swapped the HLA-E sequence with the corresponding HLA-G one downloaded from the UCSC Genome Browser in the B2M-HLA-G construct (Table S1).

We were determined to use a nonviral method for CRISPR knockin. Even if we were able to make YP001 using plasmid as the HR donor very early on, we were biased in the beginning against plasmid due to its reported toxicity despite conflicting results (*18*, *19*). Nevertheless, we confirmed successful CD19-CAR knockin in YP001 using flow cytometry and *in vitro* cytotoxicity assays (Fig 1d, S1b-d). Interestingly, both of the knockin-negative cells after two rounds of CD19-biotin sorting showed a mild yet dose-dependent cell killing response (Fig S1d), which was consistent with our detection of residual CD19-CAR^+^ cells by a G4S-linker antibody (Fig S1c). This suggests that CD19-biotin antibody may not fully sort all the target cells.

In favor of a nonplasmid nonviral knockin strategy, we settled on the closed-ended doggybone DNA (dbDNA) (*20*), which is supposed to be more stable and may therefore work better than the open-ended double-stranded DNA as the HR donor (*18*). After optimization, we were able to obtain about 2% TCRαβ^-^/CD19-CAR^+^ cells in YP001 by flow cytometry using dbDNA for the first time (Fig S1e). Importantly, the YP001ΔS control inserting a CAR lacking the CD8 signal peptide (Fig S1a) showed no knockin cells (Fig S1e); this control was very helpful in our hands to rule out false positive signals by some commercial antibodies that claim to recognize pan-single-chain variable fragment (ScFv) epitopes. After CD19-biotin sorting, the YP001 knockin yielded 2×10^6^ positive cells from a total pool of 4.5×10^7^ cells (4.4%), consistent with an approximate 4% CD19-CAR^+^ rate detected by streptavidin-FITC flow cytometry (Fig S1e). This roughly 2-fold enrichment suggested that CAR expression might be activated by antigen stimulation during sorting, and would also be consistent with incomplete sorting as described earlier (Fig S1c-d). Because 1×10^5^∼5×10^5^ CAR-T cells were used for animal studies by the Sadelain group previously (*8*, *21*, *22*), we planned to use these 2×10^6^ CD19-biotin-sorted YP001 cells for a preliminary animal study (Fig 2a).

**Fig 2.**
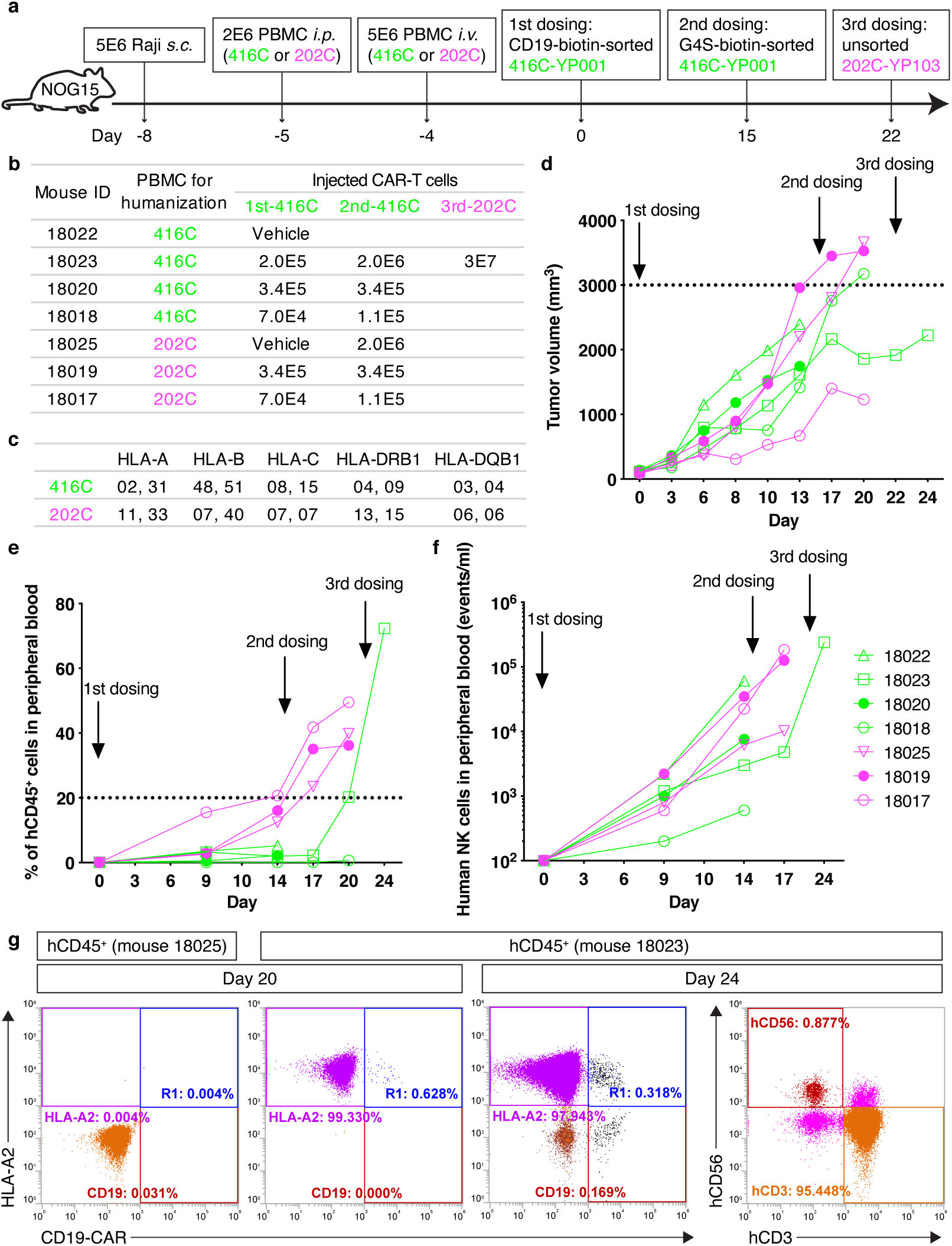
The first animal study. **a**, Timeline of the first animal study using NOG-hIL-15 (NOG15) mouse (*s.c.*, *i.p.* and *i.v.* stand for subcutaneous, intraperitoneal and intravenous, respectively). **b**, Table showing specific amount of cells used for three dosings (see **a**) in the humanized animals. **c**, Human leukocyte antigen (HLA) haplotype information for PBMC donor 416C (green) and 202C (magenta). Note that 416C is HLA-A2-positive. **d**-**f**, Data from the humanized mouse study showing tumor volume (**d**), the percent of human CD45 (hCD45) positive cells over total CD45^+^ cells (hCD45^+^ plus mCD45^+^) (**e**), and the amount of human hCD45^+^/hCD3^-^/hCD56^+^ NK cells (**f**) (see Fig S3c for flow cytometric gating strategy and percent/events conversion). A dotted line in **d** indicates that mice were supposed to be euthanized when the tumor volume was larger than 3000 mm^3^. Note that mouse 18019 was not euthanized on day 17 hoping the injected CAR-T cells might reduce tumor volume below 3000 mm^3^ after the second dosing. A dotted line in **e** indicates a 20% hCD45 reconstitution cutoff. **g**, Flow cytometric analysis for hCD45^+^ cells in peripheral blood collected from mouse 18025 and 18023 on day 20 and 24. See also Fig S2.

We decided to work with a contract research organization (CRO) that uses an innovative method by an intraperitoneal PBMC injection followed by an intravenous PBMC injection the next day into immunodeficient mice expressing human interleukin-15 (hIL-15) to further boost human T and NK cell reconstitution (Fig 2a). We chose a subcutaneous xenograft model, which is less expensive and potentially longer-lasting than a systemic model, but may be good enough to give us two readouts: a primary readout for CAR-T cell rejection by flow cytometry as long as the engrafted tumor cells can stimulate infused CAR-T cells; a secondary readout for tumor size if a tumor-inhibitory effect is strong enough.

We began with a preliminary animal study to identify a dose of CAR-T cells that could propagate in the autologous setting but be fully rejected in the allogeneic setting. To minimize PBMC-resultant GvHD, we inoculated NOG-hIL-15 (NOG15) mice (*23*) with CD19-positive Raji cells followed by prescreened PBMC (donor 416C or 202C with distant HLA haplotypes) and various doses of CD19-biotin-sorted 416C-YP001 or vehicle (Fig 2a-c). Unfortunately, we failed to detect any CAR-T cells at 9 days post CAR-T treatment (Fig S2c) without any unusual observations in the mice (Fig 2d-f, S2b and S3c). We noted that the dose previously used by the Sadelain group was for cells collected at 7 days post electroporation (*8*), and thus may not be applicable to our 2-week culture protocol (Fig 1d). We also noted the possibility that CD19-biotin-sorted cells may not function as well as naive CAR-T cells that were not pre-exposed to tumor antigens. Therefore, we switched to G4S-biotin sorting (Fig S1f) and were able to increase the dose of YP001 cells for a second dosing using the same group of animals (Fig 2a and 2b).

We also found that a Fc-tagged CD19-FITC antibody worked the best in our hands for detecting knockin cells *in vitro* among 7 antibodies we had tested (Fig S1f and data not shown), which was used in most experiments going forward.

At two days post second dosing, we were able to detect autologous CAR-T cells in peripheral blood from mouse 18023 injected with 2×10^6^ G4S-biotin-sorted YP001 cells, but not in the allogeneic control mouse 18025 where both human T and NK cells were successfully reconstituted (Fig 2e, 2f and S2d). At five days post second dosing, the immune rejection effect was more convincing when we used an HLA-A2 antibody to distinguish HLA-A2^+^ 416C-YP001 cells (Fig 2c) from reconstituted human lymphocytes in a flow cytometric analysis (Fig 2g and S2d). Notably, mouse 18023 had a flattened tumor growth curve after the second dosing (Fig 2d), suggesting an inhibitory effect. We thus confirmed both the utility of our animal model and our CAR-T dose, the latter of which is consistent with a previous report (*24*).

In the meantime, we failed to make any double knockin cells using dbDNA as the HR donor. Instead, we successfully made YP103 cells derived from PBMC 202C using plasmid as the HR donor, which were used for a third dosing of mouse 18023 humanized with PBMC 416C (Fig 2a and 2b). For this experiment we decided to use unsorted cells and test whether a humanized mouse could perform natural sorting with toleration. At two days post third dosing in mouse 18023, we were able to detect a distinct population of HLA-A2^-^/CD19-CAR^+^ cells (202C) from flow cytometry of cells (416C) that were further amplified after the second dosing (Fig 2g). The following day, we found that CD3^-^/CD19-CAR^+^ cells from 202C-YP103 (HLA-A2^-^) propagated more slowly than cells from 416C-YP001 (HLA-A2^+^) (Fig S2e). Given that the allogeneic YP001 cells were readily rejected two days post second dosing (Fig S2d), whereas the allogeneic YP103 cells were still detected three days post third dosing before mouse 18023 died (Fig S2e), our HLA-G-engineered cells might have evaded immune rejection, but it was not clear how long they could sustain.

After optimizing genome editing using plasmid HR donors, we were ready to perform another animal study to compare allogeneic YP103 with different control cells. Since the previous study revealed PBMC 202C as a better donor for mouse humanization (Fig 2e and 2f), we generated 202C-YP001, 416C-YP001 and 416C-YP103 (Fig 1d-f and S1g). Several points are worth noting: 1) Plasmid HR donors performed much better than dbDNA HR donors in terms of both editing efficiency and cell viability, possibly due to enhanced T cell activation upon sensing exogenous genomic material naturally present in bacteria (*25*); 2) The *TRAC* knockin rate was more sensitive to the T cell activation state and PBMC quality than the *B2M* knockin rate (data not shown), consistent with open chromatin-enabled editing (*26*); 3) The *B2M* knockin rate (∼30% on day 6 and 12) was higher than the *TRAC* knockin rate despite an increase on day 12 (∼10%) from day 6 (∼5%) for the latter (Fig 1f and S1g), likely consistent with a chromatin state that was less open for *TRAC* than *B2M* on day 2 when electroporation was done, followed by gradually opened chromatin states for *TRAC* during the 2-week culture; 4) The total *TRAC* knockin rate was similar between YP001 (single knockin) and YP103 (double knockin) (Fig 1e and 1f), consistent with reported coincidental HR events (*27*); 5) We failed to generate YP102ΔS on day 12 for reasons that were not clear (Fig S1g).

Since we could not use CD19-biotin for sorting and the G4S linker is present in both CD19-CAR and B2M-HLA fusion genes (Fig 1a), G4S-biotin sorting was less desirable for comparing YP001 and YP103. We therefore depleted CD3^+^ cells to minimize their impact (Fig 3a and S1h), and obtained approximately 9×10^6^ CD3^-^/HLA-ABC^-^/HLA-G^+^/CD19-CAR^+^ target cells from 416C-YP103 on day 14, sufficient for testing in 3 mice at 2×10^6^ target cells per mouse. We then calculated the required number of CD3-depleted 202C-YP001 cells that also contained 9×10^6^ target cells (CD3^-^/CD19-CAR^+^) for a side-by-side autologous/allogeneic comparison with 3 mice/group.

**Fig 3.**
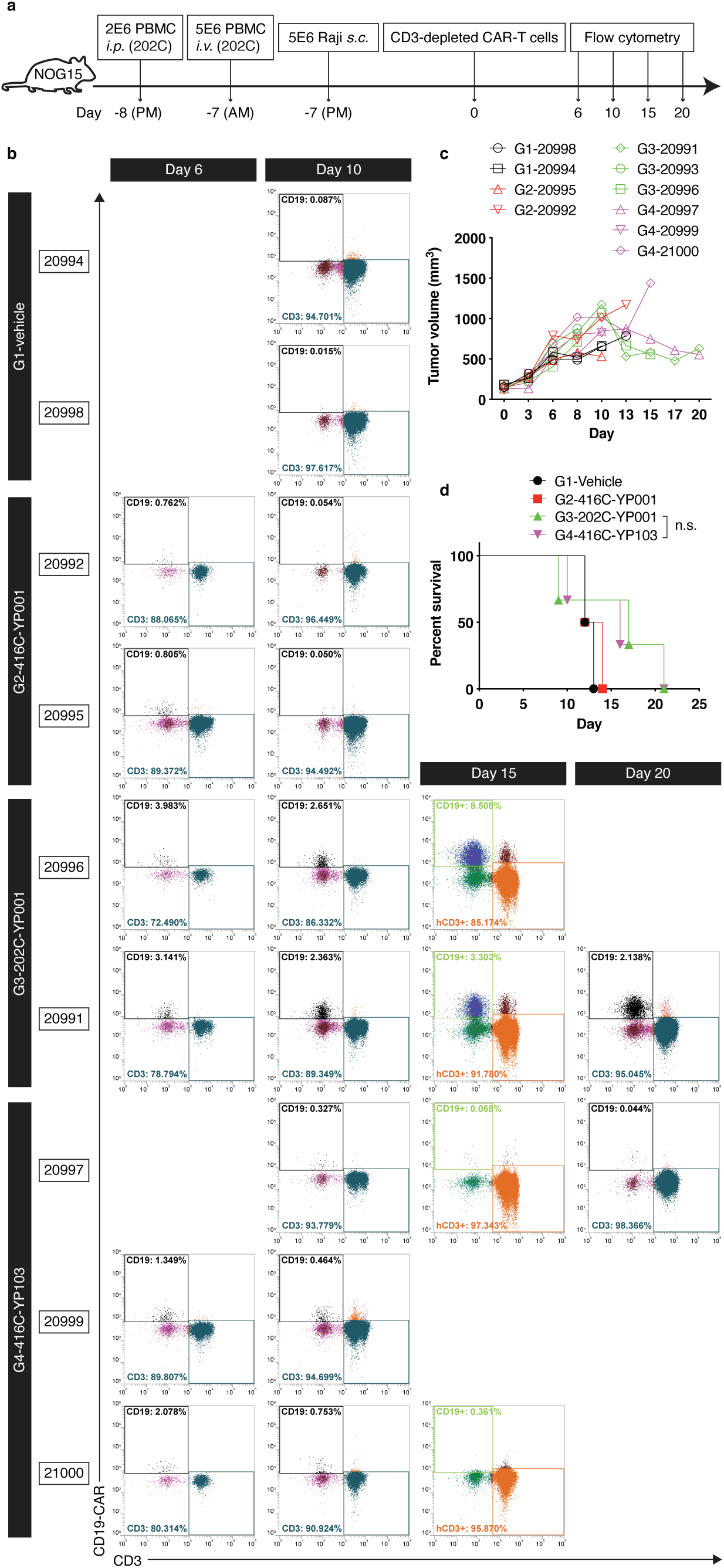
The second animal study. **a**, Timeline of the second animal study (AM and PM stand for morning and afternoon, respectively). **b**, Flow cytometric analysis for hCD45^+^ cells in peripheral blood collected from selected 6 animals out of total 10 animals on day 6, and all the surviving animals on day 10, 15 and 20. **c**, Tumor volume for individual animals. **d**, Survival curve for 4 groups with n.s. indicating a statistically insignificant difference between the G3 and G4 groups (*P* value = 0.95 using the log-rank Mantel-Cox test). Note that mouse G2-20992 received a second dosing on day 13 but died one day later, and mouse G1-20998 was euthanized due to a 20% plus body weight loss that was not recovered for 72 hours. See also Fig S3.

In our initial study, the hCD45 leukocyte rates were too low for grouping animals (Fig S2a), so we injected 202C-PBMC before Raji in 10 mice (Fig 3a) and were able to group them based on both tumor volume and hCD45% (Fig S3a). We selected 6 mice for 202C-YP001 and 416C-YP103 testing (3 randomized mice/group), using the remaining 4 mice for vehicle and 416C-YP001 testing (2 mice/group upon randomization). Because knocking out *B2M* removes surface expression of HLA-ABC including HLA-A2, we could not utilize the HLA-A2 antibody to distinguish 416C-YP103 from host lymphocytes derived from 202C.

All injected CD3^-^/CD19-CAR^+^ cells could be detected by flow cytometry at 6 days post treatment when the mice were likely not fully humanized (Fig 3a, 3b and S3b). On day 10, the autologous 202C-YP001 and allogeneic 416C-YP103 cells could still be detected, but the allogeneic 416C-YP001 control cells were only detectable at a level similar to the vehicle controls suggesting that they were immune rejected (Fig 3b). On day 15, 416C-YP103-injected mouse 21000 appeared to have lost CAR-T expression (Fig 3b), correlated with abruptly increased tumor volume (Fig 3c) and followed by death on day 16 (Fig 3d). However, in mouse 20997, another 416C-YP103-injected animal, CAR-T cells were detected continuously until day 20 (Fig 3b). Since all the cells and animals were isogenic, the differential phenotypes between the two mice might reflect a dose response caused by cell heterogeneity. Injected CD3-depleted 416C-YP103 mix contained ∼5% HLA-ABC^-^/HLA-G^+^/CD19-CAR^+^ target cells that were expected to evade immune rejection, and ∼3% HLA-ABC^-^/HLA-G^-^/CD19-CAR^+^ cells that were expected to be rejected by activated NK cells (Fig 1f). Therefore, there is a possibility that an uneven cell distribution between the 3 mice led to more HLA-ABC^-^/HLA-G^-^/CD19-CAR^+^ cells with a possible delayed immune rejection by NK cells, and fewer target cells that may not be enough to win over Raji cells in mouse 21000, leading to tumor relapse (Fig 3b and 3c). To minimize heterogeneity, we were able to enrich our target cells from YP103 by G4S-biotin sorting in a separate experiment (Fig S1i).

Notably, autologous 202C-YP001 cells amplified at a drastically higher rate than allogeneic 416C-YP103 cells (Fig 3b), despite similar survival curves for the two groups (Fig 3d) and similar controlled tumor sizes in mice 20991 and 20997 (Fig 3c). This is reminiscent of the previously observed slower growth of 202C-YP103 compared to 416C-YP001 (Fig S2e), and suggests a rate-limiting mechanism that may be related to HLA-G engineering. If this observation holds true in clinics, HLA-G-engineered CAR-T cells may have a safety benefit from reduced cytokine release syndrome (CRS) risk.

Compared to our first animal study, the mice in our second study tended to die much earlier with smaller tumor sizes (Fig 2d and 3c), which was likely due to increased GvHD imposed on the mice and non-specific anti-tumor effects caused by injection of PBMC before Raji cells (*28*). Ironically, by the time when the CRO received our newly prepared YP100 (Fig S1j), there was only one surviving animal without CAR expression (mouse 20992) that could be used for a second dosing on day 13, which however died one day later. Since recent work from the Sadelain group showed *B2M*-deficient CAR-T cells are immune rejected by NK cells in a humanized mouse model (*21*), we believe our allogeneic YP100 cells would behave similarly.

Finally, we worked with a contract development and manufacturing organization (CDMO), where YP103 was successfully reproduced after a technique transfer. We have thus reprogramed CAR-T cells with an immune privilege by targeting an HLA-G single-chain trimer to the *B2M* locus using a cost-effective plasmid-based CRISPR editing method. HLA-G engineering is poised to unleashing genome editing and iPSC technologies in clinics after removing the immune barrier.

## Acknowledgements

We thank every single individual who has helped along this endeavor.

## Funding

This work was supported by Yuanpei Therapeutics.

## Author contributions

Y.X. conceived the project, provided the funding, led all the work, and wrote the manuscript.

S.H. did all the *in vitro* work. The CRO did all the *in vivo* work, the identity of whom cannot be disclosed in compliance with confidentiality clauses. Y.X., S.H. and C.X. cloned the constructs.

## Competing financial interests

The authors declare no competing financial interests.

## Materials and methods

### Motif analysis

A 73 bp reference DNA sequence spanning the 23 bp B2M gRNA sequence (20 bp spacer plus 3 bp PAM) with 25 bp sequences extended from each end was downloaded from the UCSC Genome Browser (hg38 assembly). The other two 73 bp sequences with mutations illustrated in Fig 1b for Yuanpei (our study) and CN110616189B, along with the reference sequence, were used for motif analyses by FIMO (version 5.5.7) from the MEME Suite with a default 1×10^-4^ *P* value cutoff against the JASPAR motif database (2024_CORE_vertebrates_non-redundant).

### Homologous recombination donor

The 1 kb homology arms for *TRAC* and *B2M* were PCR amplified from genomic DNA isolated from PBMC of one healthy donor, except that the right homology arm for *B2M* was synthesized after multiple failed PCR attempts. Fusion genes including homology arms (Table S1 and Fig S1a) were cloned into a pUC57-GW-Amp vector (GENEWIZ) connected with an XbaI site (adjacent to the vector) and a TelRL (*20*) site (adjacent to the insert) in both ends, with glycerol stocks stored at-80 °C. Extracted plasmids using a commercial kit (TIANGEN, DP120) were either used directly as the HR donor (eluted with nuclease-free water followed by concentration in a heat block with lid open at 70 °C until 2.5 µg/µl or more was reached), or used as templates to generate dbDNA (*20*). Briefly, plasmids were processed with TelN digestion first to generate two closed-ended DNA, followed by sequential XbaI and T5 exonuclease digestions to remove the vector, and PEG purification (*30*).

### Genome editing

CD4^+^ and CD8^+^ T cells were sorted at a 1:1 ratio in the running buffer (PBS, pH = 7.2, 0.5% BSA, 2 mM EDTA; precooled freshly-made buffer after sterilization via a 0.22 µm filter) from thawed PBMC (MileCell Bio, 10 or 25 million cells) using magnetic selection (GenScript, L00863 and L00864) in column (GenScript, CytoSinct gM columns; or XinBio, SophMag xL columns), and activated for 48 hours (37 °C, 5% CO_2_) with CD3 and CD28 stimulation (Miltenyi, TransAct, 130-128-758; or T&L Biotechnology, ActSep, GMP-TL603-1000) in the T cell stimulation media (STEMCELL, ImmunoCult-XF T Cell Expansion Medium, 10981; supplemented with fresh aliquoted hIL-2 at 300 IU/ml) at a density of 1E6 cells/ml per manufacturer’s instructions. On day 2 (Fig 1d), 1.25E6 activated T cells were prepared in 25 µl total volume with electroporation buffer (Gibco, Opti-MEM, 31985070) to maintain a cell density at 5E7 cells/ml for electroporation with 1.2 µM SpCas9 (Kactus Biosystem, CAS-EE109) and 1.8 µM sgRNA (GENEWIZ; DSL grade) that were preincubated for 10 minutes at room temperature, and 4 µg plasmid per gene targeting, in a 20 µl cuvette using the Celetrix electroporation system (CTX-1500A LE+, “Cell Line” program, 540 V, 20 ms). In the case of 100 µl cuvette electroporation, 5.5E6 cells were prepared in 110 µl electroporation buffer with 20 µg plasmid. In the case of dbDNA as the HR donor, 2 µg or 10 µg dbDNA was used in the 20 µl or 100 µl cuvette, respectively. In the case of double knockin to make YP103, 2.4 µM SpCas9, 1.8 µM TRAC sgRNA, 1.8 µM B2M sgRNA, 4 µg YP1 plasmid and 4 µg YP3 plasmid were used (Fig 1d). After electroporation, cells were let sit in the cuvette for 15 minutes at room temperature before transferring to the plate with prewarmed T cell stimulation media prepared at a density of 1E6 cells/ml (37 °C, 5% CO_2_). On day 5, based on the color of the media, a certain amount of the T cell stimulation media was added to the cells accordingly. On day 6, 3E5 cells were collected for flow cytometry (Beckman Coulter, CytoFLEX S) with selective antibodies used at a 1:200 dilution except as noted: Zombie-BV421 (Biolegend, 423113, 1:400), TCRαβ-APC (Biolegend, 306718), CD4-Percp-cy5.5 (Vdo Biotech, S0028), CD8-PE (Vdo Biotech, S0044), CD3-APC (Vdo Biotech, S0017), HLA-ABC-AF700 (Biolegend, 311438), HLA-E-pE-cy7 (Biolegend, 342608), HLA-G-pE-cy7 (Biolegend, 335911), G4S-PE (Hycells, GS-ARPE25), FMC63-PE (ACROBiosystems, FM3-HPY53), CD19-FITC (Fc-tagged; ACROBiosystems, CD9-HF251, 1:40). Note that CD8-PE was sometimes omitted due to antibody conflict, and in such a case, only CD4-Percp-cy5.5 was used to gate CD4^+^ and CD8^+^ T cells (Fig S1b). T cells were then maintained at a density of 1E6 cells/ml by adding T cell expansion media (STEMCELL, ImmunoCult-XF T Cell Expansion Medium, 10981; or ExCell Bio, OptiVitro UniEx T Cell Serum-free Medium, TE000-N052) including 300 IU/ml hIL-2 every 2 days. On day 12 or day 13, another flow cytometric analysis was done to confirm genome-edited cells. On day 14, cells were either positively selected in the running buffer using CD19-biotin (ACROBiosystems, CD9-H8259, 1:10) or G4S-biotin (Hycells, GS-ARBI100, 10 µg/ml), followed by streptavidin bead (Vdo Biotech, CSM0150SA-2, 10 µl beads per 1E7 cells) incubation (or streptavidin-FITC staining for flow cytometry using #405201 antibody from Biolegend), or negatively selected in the running buffer by CD3 nanobeads (GenScript, L00896, 6 µl nanobeads per 1E7 cells), per manufacturer’s instructions except as noted. Sorted cells were resuspended in freezing media (MileCell Bio, Kryogene Cell Freezing Media-CGT, AR0008-100) and stored in a freezing container (Kemesser Technology, CELLHOME-12) at-80 °C for one overnight before transfer to liquid nitrogen.

### sgRNA sequence

TRAC (guide: CAGGGTTCTGGATATCTGT): C*A*G*GGUUCUGGAUAUCUGUGUUUUAGAGCUAGAAAUAGCAAGUUAAAAUAAG GCUAGUCCGUUAUCAACUUGAAAAAGUGGCACCGAGUCGGUGCU*U*U*U B2M (guide: GGCCACGGAGCGAGACATCT): G*G*C*CACGGAGCGAGACAUCUGUUUUAGAGCUAGAAAUAGCAAGUUAAAAUAA GGCUAGUCCGUUAUCAACUUGAAAAAGUGGCACCGAGUCGGUGCU*U*U*U Asterisk (*) denotes 2′-O-methyl 3′ phosphorothioate

### *In vitro* cytotoxicity assay

CAR-T cells were cocultured with luciferase-expressing Raji cells (provided by the CRO) in duplicate at the indicated effector (E) to target (T) ratios (Fig S1d) in a 96-well plate (Thermo Fisher, Nun MicroWel, 136101) with 1E5 target cells in total 100 µl RPMI 1640 media (Gibco, C11875500BT) including 2% fetal bovine serum (FBS, ExCell Bio, FSP500) per well. Target cells alone were plated to determine the maximal luciferase expression in relative light units, or RLUmax. Eighteen hours later, 100 µl luciferase substrate (VKEY Bio, D-luciferin, A2001001N) was added to each well followed by plate reader measurement (PerkinElmer, VICTOR Nivo 5S). Cytotoxic lysis was determined as (1 − RLUsample/RLUmax) × 100%.

### Humanized mouse model

6-8 week old female NOD.Cg-*Prkdc^scid^IL2rg^tm1Sug^*Tg(CMV-IL2/IL15)1-1Jic/JicCrl mice, or NOG-hIL-15 (Charles River, Beijing) in short, were allowed to acclimate to the animal facility at the CRO for 5-7 days in a group-housing setting with 2–3 mice per cage in a reverse 12 hour light/dark cycle with ad libitum access to food and water. The experimental protocols were approved by the Institutional Animal Care and Use Committee (IACUC) via the CRO. Mice were inoculated with Raji cells, PBMC and CAR-T cells followed by flow cytometric analyses with peripheral blood at indicated time as shown in Fig 2a and Fig 3a. Specifically, Raji cells were cultured in Raji media (RPMI 1640 media including 10% FBS) before subcutaneous engraftment in the right flank of the mouse in a 100 µl solution including 5E6 cells in PBS and Matrigel mixed at a 2:1 volume ratio. PBMC from donor 416C or 202C (MileCell Bio) were thawed in Raji media and resuspended in 200 µl PBS before intraperitoneal or intravenous injection into the mouse. CAR-T cells or vehicle (freezing media) were intravenously injected into the mouse at a volume of up to 250 µl immediately upon thawing; a small number of cells was used for cell counting. Body weight and tumor volume (TV = 0.5 × a × b^2^) were measured 2-3 times per week, with the length and width of the tumor obtained using calipers as a and b, respectively. Antibodies used for flow cytometric analysis include mCD45 (Alexa Flour 700; Thermo Fisher, 56-0451-82), hCD45 (APC-eFluor 780; Thermo Fisher, 47-0459-42), hCD56 (PE; Thermo Fisher, 12-0567-42), hCD3 (Pacific Blue; Biolegend, 300431) with m standing for mouse and h standing for human. Two FITC-labeled CD19-CAR antibodies were used (ACROBiosystems CD9-HF2H2 for day 6 and day 10 in Fig 3b; ACROBiosystems CD9-HF251 for all the remaining data). The percent of hCD45^+^ cells was calculated by equation hCD45^+^/(hCD45^+^ + mCD45^+^) × 100%. hCD45^+^/hCD3^-^/hCD56^+^ cells were counted as human NK cells. Mice were euthanized by CO_2_ when encountering a 20% plus body weight loss that was not recovered for 72 hours or when the tumor volume was larger than 3000 mm^3^, or on other conditions specified by the IACUC.

## Statistical analysis

No statistical methods were used to predetermine sample size. A log-rank Mantel-Cox test was performed for statistical analysis in Fig 3d using the GraphPad Prism software (version 9).

---

**Fig S1.**
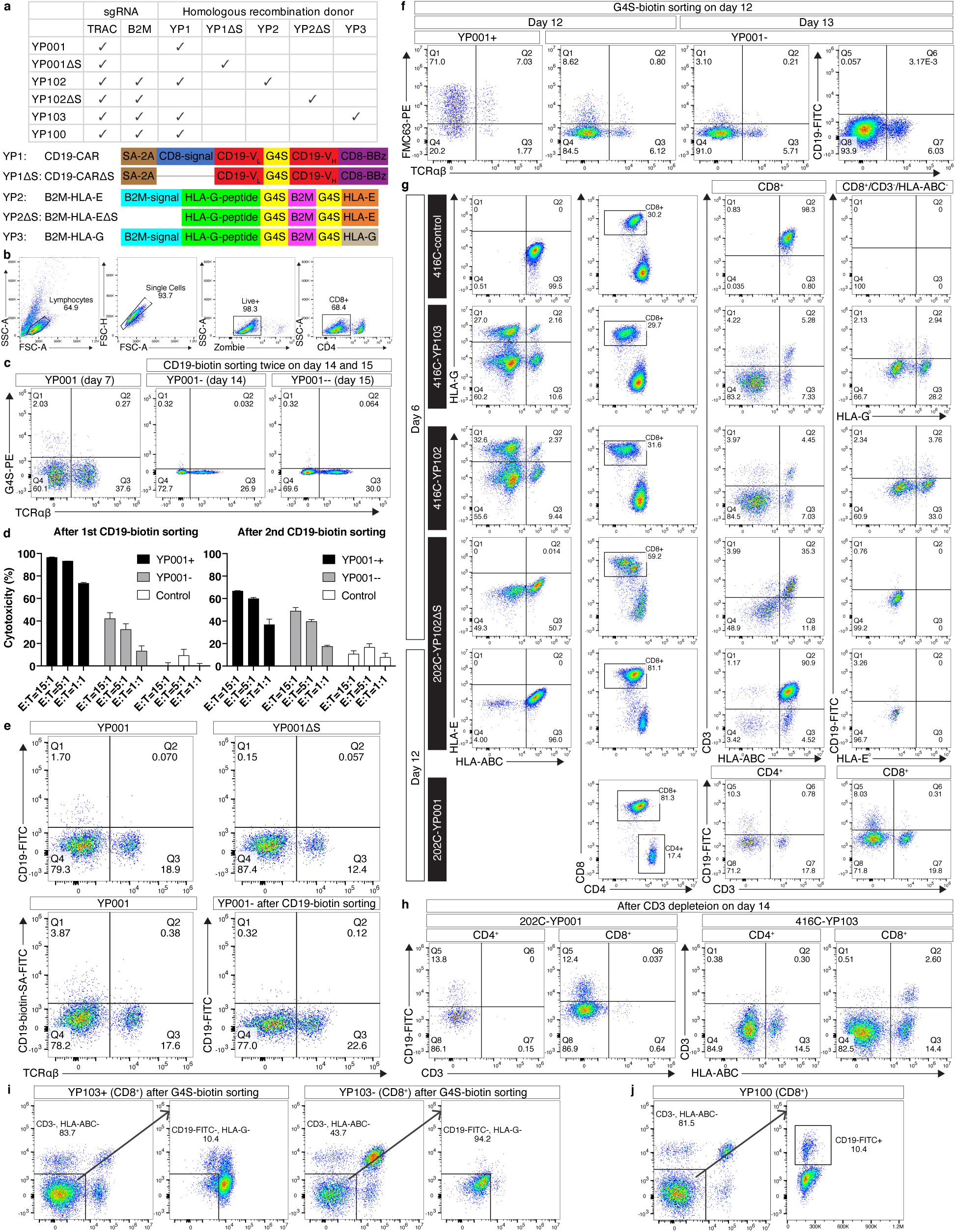
Additional *in vitro* data. **a**, Genome editing materials for generating different CAR-T cells (upper), with major structural components illustrated in the fusion gene schematic (lower). Note that YP001ΔS and YP002ΔS used constructs lacking the CD8 signal peptide (CD8-signal) and B2M signal peptide (B2M-signal), respectively. See also Table S1. **b**-**d**, Flow cytometry (**b**, **c**) and *in vitro* cytotoxicity assay (**d**) after two rounds of CD19-biotin sorting for YP001. Flow cytometric gating strategy (**b**) and knockin result (**c**, left) for YP001 at 7 days post thawing (day 7, see Fig 1d) are shown. Data include positive (YP001+) and negative (YP001-) cells after the first sorting on day 14, the latter of which were subject to another round of sorting on day 15 leading to positive (YP001-+) and negative (YP001--) populations. Three effector (E) to target (T) ratios were used with unedited cells as controls running two technical replicates (data are mean ± SD) in **d**. **e**, Flow cytometric analysis for CD8^+^ YP001 before and after sorting with CD19-biotin on day 13, with YP001ΔS as a control. Note that YP001 had increased CD19-CAR^+^ cells after sorting using a FITC-labeled streptavidin (SA) antibody (lower left). Also note increased TCRαβ^+^/CD19-CAR^-^ cells in YP001- cells after sorting (lower right) than before sorting (upper left). **f**, Flow cytometric analysis for CD8^+^ YP001 sorted with G4S-biotin on day 12. Note non-specific FMC63-PE signals in YP001- (negative cells post sorting) that were absent by CD19-FITC staining. **g**, **h**, Flow cytometric analysis for various cells on day 6 and 12 (**g**), and day 14 after CD3 depletion (**h**). Note incomplete CD3 depletion in the CD8^+^ YP103 (**h**). **i**, **j**, Flow cytometric data of positive (YP103+) and negative (YP103-) cells after a G4S-biotin sorting (**i**), and successful generation of YP100 (**j**). Closed-ended doggybone DNA (dbDNA) was used as the HR donor for data in **e** and **f**, with the rest using plasmid as the HR donor. CD4^+^ and CD8^+^ T cells were sorted from thawed PBMC before activation in all experiments (Fig 1d), with flow cytometric data sometimes just showing the CD8^+^ gate to minimize data redundancy.

**Fig S2.**
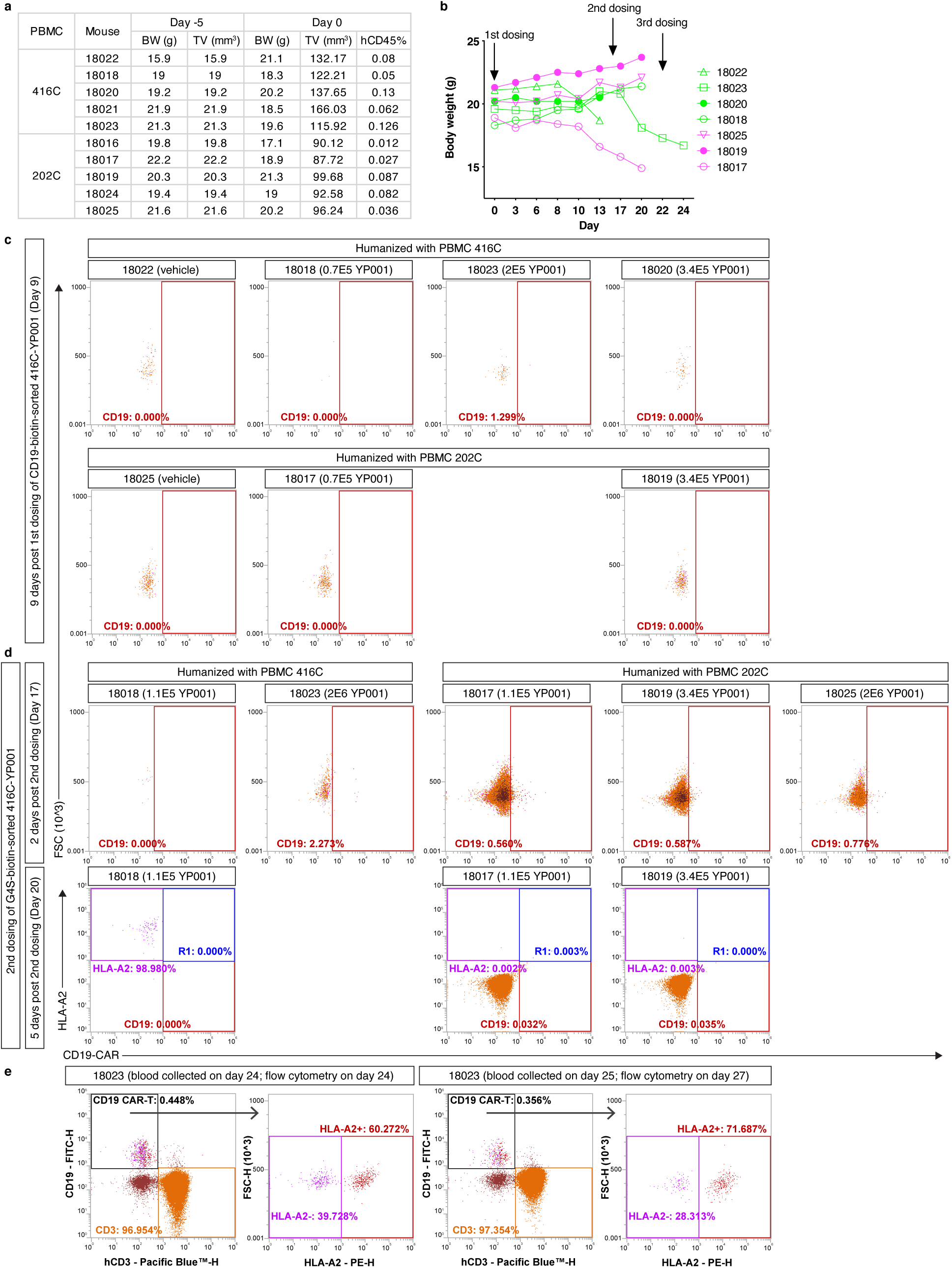
Additional data for the first animal study. **a**, body weight (BW), tumor volume (TV) and hCD45% data on day-5 and 0 (see Fig 2a). BW and TV on day-5 were used for randomization and grouping for injection with PBMC 416C or 202C. **b**, Body weight of all the animals throughout the experiment. **c**-**e**, Flow cytometric analysis for hCD45^+^ cells in peripheral blood collected from different animals. Peripheral blood collected from mouse 18023 on day 25 was stored at 4 °C until day 27 when flow cytometry was done.

**Fig S3.**
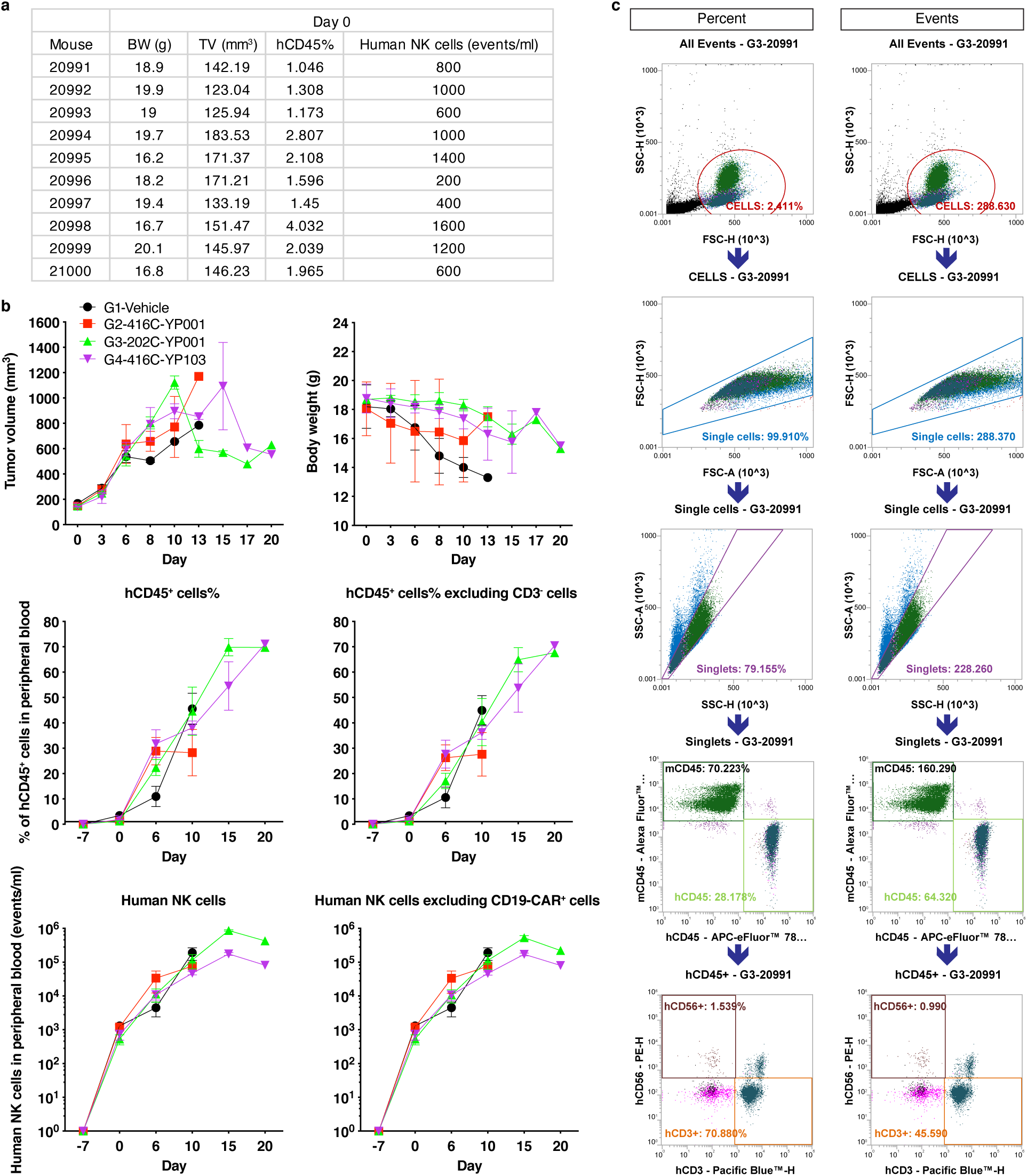
Additional data for the second animal study. **a**, **b**, Animal information on day 0 used for grouping before CAR-T treatment (**a**), and throughout the experiment (**b**). Because CAR-T cells were depleted with CD3^+^ cells without a G4S-biotin or CD19-biotin sorting, CD3^-^ cells that were potentially introduced from CAR-T treatment were excluded from total hCD45^+^ cells in a separate analysis (middle right). We noticed hCD56^+^/CD19-CAR^+^ cells, which were also excluded in a separate analysis (lower right). **c**, Flow cytometric gating strategy by the contract research organization (CRO) for mouse 20991 on day 6 (see Fig 3a) in either percent (left) or events (right). Data are mean ± SEM in **b**.

**Table S1.**
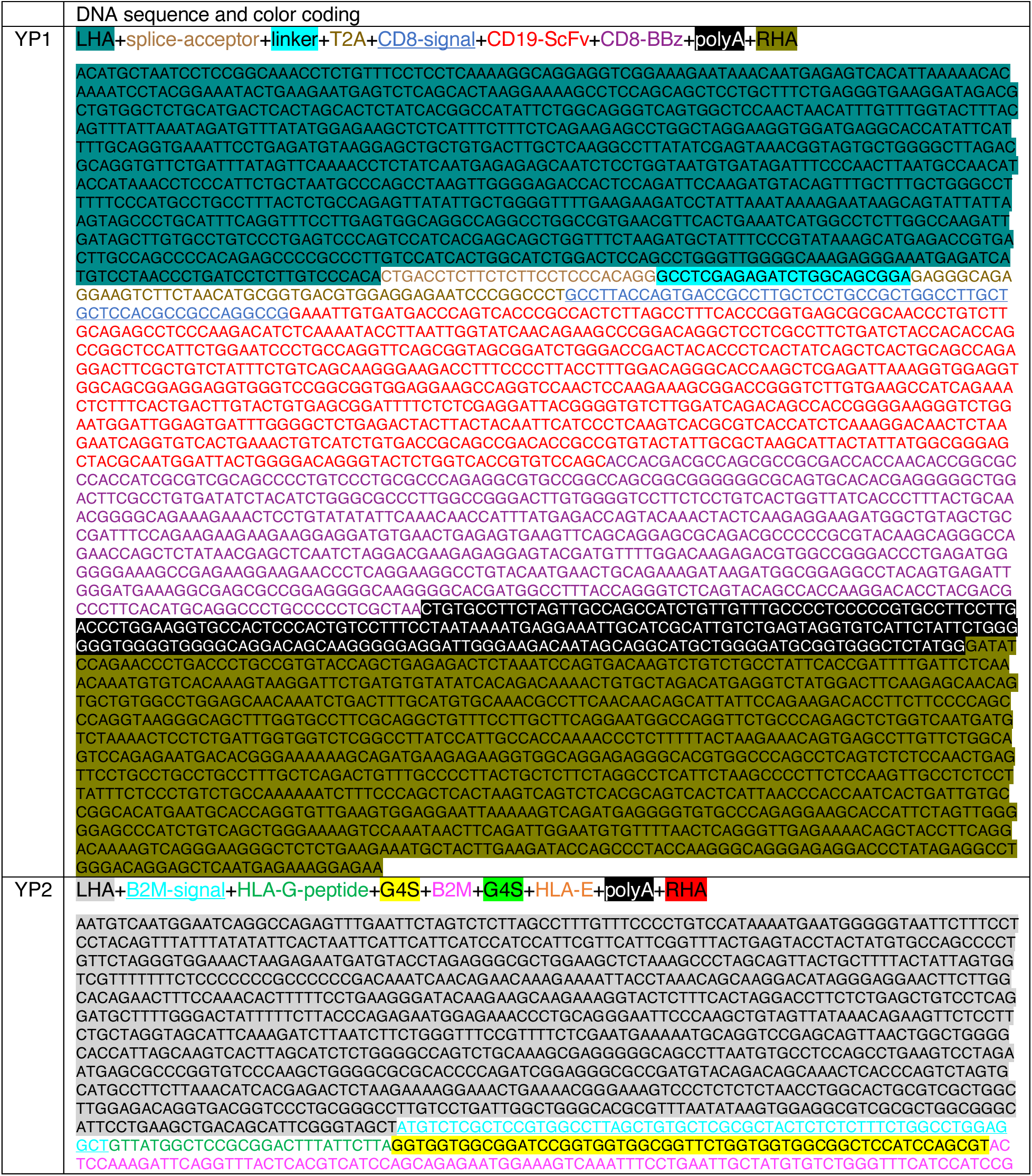
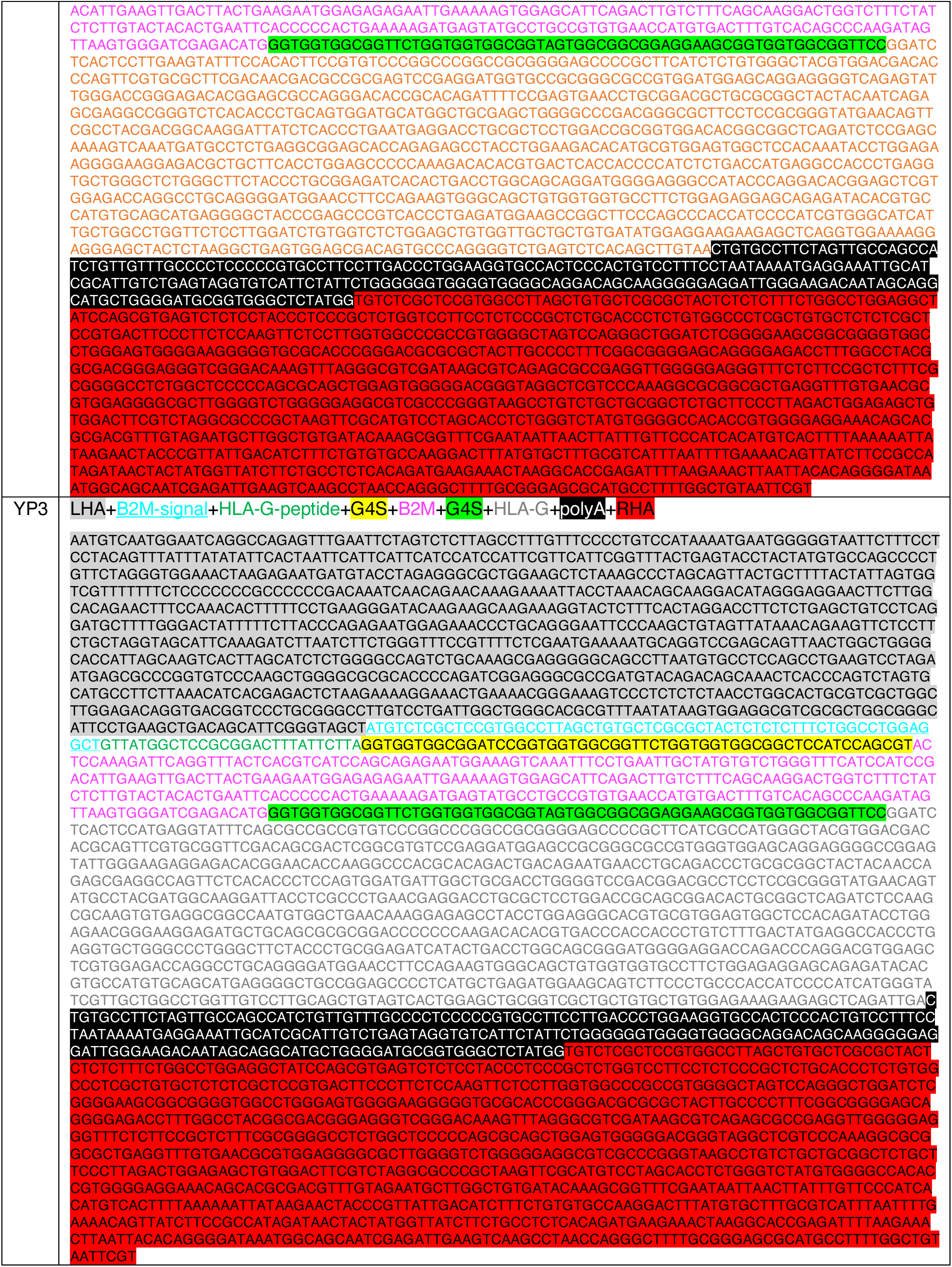
DNA sequences of the main constructs. The sequences for CD19-ScFv and CD8-BBz (including the hinge region, transmembrane domain, and etc.) are from the patent CN107793480B. The sequences for splice-acceptor (SA), linker, T2A (a self-cleaving 2A sequence) and polyA are from https://www.ncbi.nlm.nih.gov/nuccore/2633523300.

## Notes

### Competing Interest Statement

The authors have declared no competing interest.

